# Clarifying misconceptions of biomolecular condensate formation

**DOI:** 10.1101/2024.07.17.603945

**Authors:** Nidhi Hosamane, McKayla Hartman, J. Matthew Dubach

## Abstract

Biomolecular condensates are liquid-like phase separations of disordered proteins and other molecules within cells. These membrane-less organelles are essential in critical cellular processes and are aberrant in myriad disease. Yet, the thermodynamics of condensate formation are not entirely understood, thus the mechanisms of condensate function remain elusive. Two assumptions about biomolecular condensates are that they form through liquid-liquid phase separation (LLPS) and require multivalent interactions. Here, we show results and propose thermodynamic frameworks that suggest these assumptions are not always correct. We demonstrate that condensation does not follow canonical LLPS and that liquid-like phase separation can arise from monovalent interactions. These results clarify the mechanism of condensate thermodynamics to reframe our understanding of condensates in cellular function and disease.

## Introduction

Biomolecular condensates^1^ were first described as a liquid-like phase separation 15 years ago^2^. While separated bodies such as the nucleolus^3^ had been well established, the ramifications of the physics governing liquid-like phase separation and the biological impact generated profound interest across biology^4^. Phase separation has been associated with critical cellular functions, such as transcription^5^, and myriad disease^6^, such as neurodegeneration^7^. Further investigation and understanding of condensates, the physics driving their existence, and the biological impact of their formation could reframe our understanding of cell biology.

Unfortunately, the mechanism of biomolecular condensate formation has remained elusive. Existing theories are based on polymer physics and classic thermodynamics^8,9^, yet molecular modeling and simulations are necessary to best capture the behavior of phase separating molecules in the complex cellular environment^10^. However, the current theories fail to completely describe the physics of phase separation^11^, which demonstrates that at present our understanding of the mechanism driving condensate formation remains incomplete. Instead of our nascent knowledge of complex network interactions preventing a complete thermodynamic explanation of condensates, we propose there are actually two key incorrect assumptions about condensation that are limiting the models – condensates are liquid-liquid phase separations (LLPS)^12^ and interactions within a condensate must be multivalent^11^.

Under LLPS thermodynamics a saturation concentration (C_sat_) exists above which the system forms liquid-liquid phase separation (**Fig. 1A**)^12^. When the total concentration reaches the dense phase concentration (C_den_) phase separation is no longer present and the system exists at a uniform dense phase concentration. LLPS dictates that between C_sat_ and C_den_ phase separation is the energetic minimum state producing a high concentration dense phase and a low concentration dilute phase. In LLPS the concentrations of these two phases are constant at all total protein concentrations where phase separation occurs. Increasing the concentration above C_sat_ will increase the volume fraction of the dense phase but not impact the partition coefficient (the concentration ratio of the dense and dilute phases). Thus, in the presence of phase separation the partition coefficient is independent of the total concentration.

**Figure 1:**
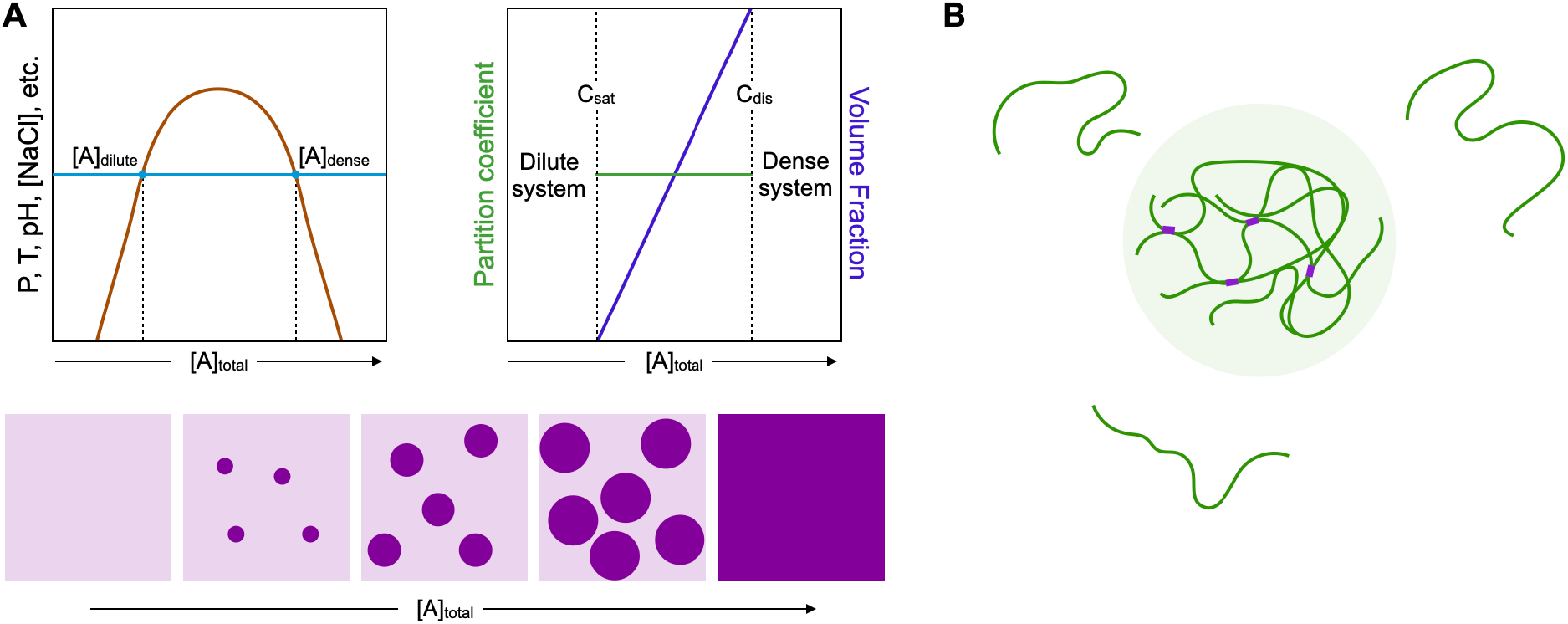
Assumed behavior of biomolecular condensates. **A)** Biomolecular condensates are thought to follow LLPS thermodynamics, generating a phase separation diagram dependent on total protein concentration and another system parameter, such as pressure or temperature. Within a constant system in a regime where phase separation occurs, increasing a concentration tie line can be drawn to determine the dilute phase and dense phase concentrations upon phase separation. Under LLPS, at all concentrations where phase separation occurs, the dilute phase concentration and dense phase concentration are constant, producing a concentration independent partition coefficient. The volume fraction of the dense phase is concentration dependent. **B)** Multiple copies of a disordered protein undergo phase separation. Within the dense phase there are numerous multivalent interactions, which provides the mechanism of biomolecular condensate formation.

Multivalency is also assumed to be necessary for biomolecular condensate formation^13^. Long disordered proteins can have numerous interactions with multiple other molecules (**Fig. 1B**). Molecular simulations show that multivalent interactions enable condensates to be stable in equilibrium with the dilute phase. However, the complexity of multivalent interactions is difficult to completely model and, since we currently do not understand the mechanism of biological phase separation, multivalency is thought to be essential to fill this gap in our knowledge. Thus, it is currently assumed that monovalent interactions cannot form condensates.

Here we show that biomolecular condensates do not follow canonical LLPS rules and that liquid-like phase separation can occur through monovalent interactions. We expect that reconsideration of condensate thermodynamics without the constraints of incorrect limitations will be critical to solving the mechanisms of condensate formation and clarifying their biological impact.

## Results

### Evidence that Biomolecular condensates are not LLPS

LLPS dictates that under phase separation the partition coefficient is constant at all total protein concentrations^11^. Therefore, if LLPS was an accurate model of biomolecular condensates in cells we would expect the partition coefficient to be independent of total protein expression. To test this assumption, we measured condensate partition coefficients in three well established cellular condensates across four cell lines. Here, we used immunofluorescence on fixed cells, which allowed us to measure endogenous protein phase separation. We found that all three condensates in each cell line displayed concentration dependent partition coefficients (**Fig. 2A-C and S1**). The distribution of partition coefficient vs. total immunofluorescence intensity in single cells was different between cell lines. For example, PML condensates in MCF7 cells were limited to lower levels and highly dynamic partition coefficient regions while PML in HCC1937 cells was at higher levels with lower partition coefficient variance. Other markers, such as PML in OV90 cells, showed a wide range of overall intensity vs. partition coefficient. Strikingly, combining single cell data from all cell lines for each marker generated a consistent non-linear relationship between partition coefficient and total expression. Thus, despite differences in marker expression levels, and most likely unequal levels of other protein expression between cell lines, the partition coefficient has a well-defined dependency on overall cellular expression.

**Figure 2:**
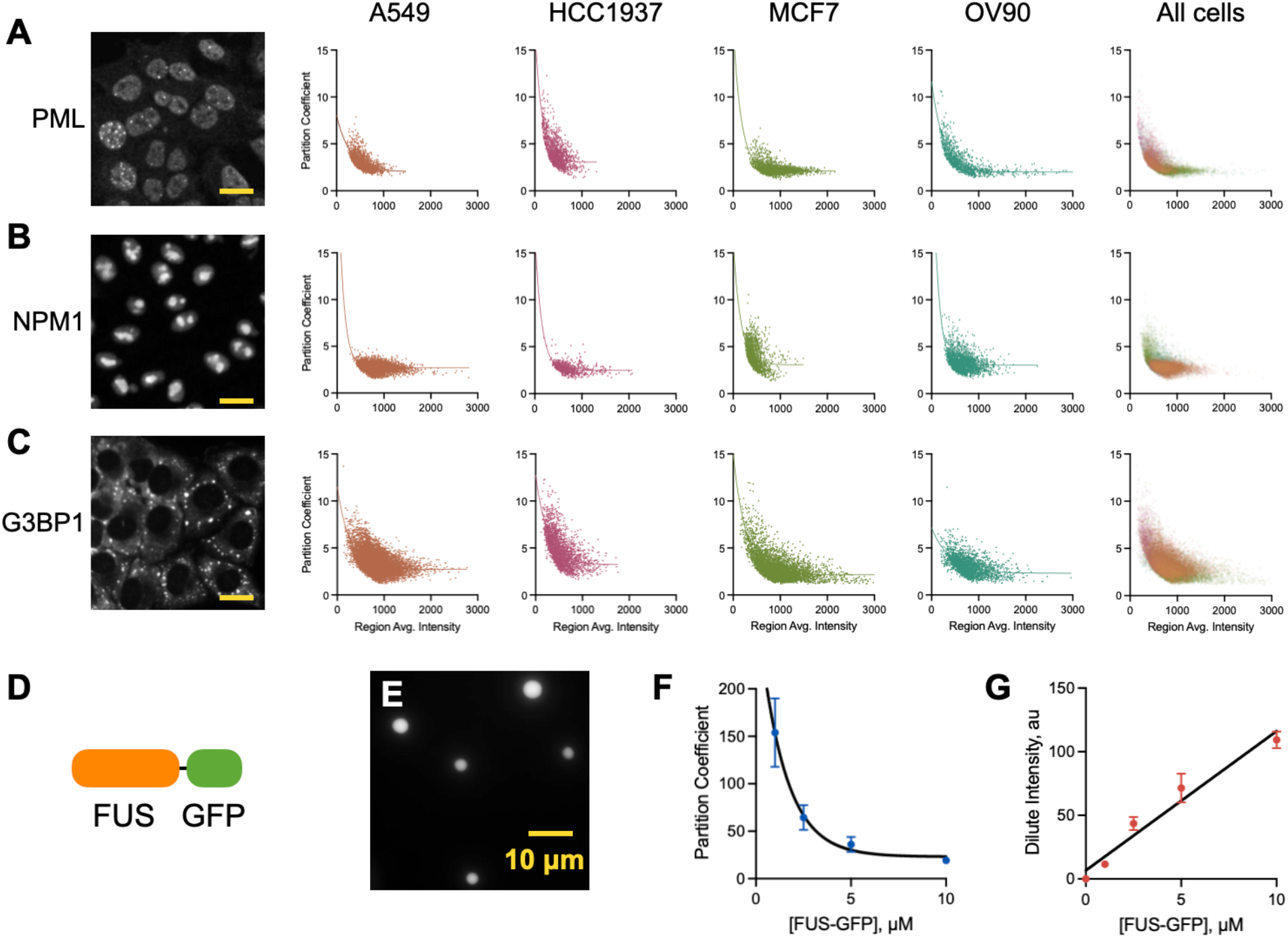
Biomolecular condensates have a concentration dependent partition coefficient. **A-C)** Representative image of PML (**A**), NPM1 (**B**) and G3BP1 (**C**) in A549 cells and the corresponding partition coefficient between condensates and the dilute phase as a function of total expression for four different cell lines, and all cell lines combined. Measurements of PML and NPM1 were performed in segmented nuclei while measurements of G3BP1 were performed in segmented cytoplasm. Shown are single cell measurements with one phase decay, ≥ 14 biological repeats and ≥ 818 cells per condition. Scale bars = 20 μm. **D)** Cartoon of recombinant FUS tagged with GFP. **E)** Representative fluorescent image of FUS-GFP forming biomolecular condensates. **F)** FUS-GFP condensate partition coefficient as a function of total FUS-GFP concentration in the system, shown are average with standard deviation, n ≥ 6. **G)** The dilute phase intensity as a function of total FUS-GFP concentration in the system, shown are average with standard deviation, n = 6.

The observation that the partition coefficient is concentration dependent in cells has been previously made by Riback, *et al*^14^. The authors calculate the standard Gibbs free energy of transfer as 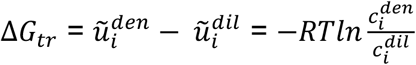. Where, the middle term is the difference in the standard chemical potential of existing in each phase (dense and dilute) and the term on the right, a function of the concentrations (*c*) in each phase, is the definition of the entropic cost of de-mixing at 50% volume fraction (the energy required to keep a molecule in the dense phase from diffusing into the dilute phase). Unfortunately, Riback, *et al* did not consider the condensate volume fraction in these calculations. However, Riback, *et al* conclude that the concentration dependent partition coefficient is driven by multicomponent interactions within condensates. Therefore, an increasing concentration of one component creates more homotypic interactions that are less energetically favorable than heterotypic interactions and destabilizing (less negative Δ*G*_*tr*_) to the condensate, reducing the partition coefficient in a concentration dependent manner. Yet, curiously, the authors show that *in vitro* condensates formed by NPM1 alone have a more negative Δ*G*_*tr*_ than a multicomponent system including NPM1, implying that homotypic interactions are actually energetically favorable (**Fig. S2**). Furthermore, why increasing the concentration of NPM1 in a multicomponent condensate, thus generating more homotypic interactions, causes Δ*G*_*tr*_ to diverge from NPM1-only condensate Δ*G*_*tr*_ remains unclear. If heterotypic interactions were drivers of a concentration dependent partition coefficient, we would expect convergent behavior as a function of total NPM1 concentration (**Fig. S2**). Ultimately, the conclusions that multicomponent interactions drive a concentration dependent partition coefficient rely upon single component condensates generating concentration independent partition coefficients as LLPS dictates. Yet, the single component NPM1 condensates measured in Riback, *et al* included PEG as a molecular crowder. PEG has recently been shown to cause NPM1 condensates to form highly viscous gels that are not in equilibrium with the dilute phase^15^. Therefore, to determine if single component condensates are indeed independent of total concentration, we measured the partition coefficient of FUS tagged with GFP - a disordered protein that can form condensates *in vitro* in the absence of molecular crowding^16^. Here, we found that the partition coefficient of FUS-GFP condensation was indeed dependent on total concentration (**Fig. 2D-F**). The dilute phase concentration was also dependent on the total FUS-GFP concentration (**Fig. 2G**). These quantitative measurements are similar to previous results that qualitatively appear to have a total concentration dependent dilute phase concentration in single component systems^16^. Therefore, single component condensates can also have a concentration dependent partition coefficient demonstrating that heterotypic interactions are not the cause of concentration dependence and, subsequently, condensate formation is likely not canonical LLPS.

### Evidence that biomolecular condensates do not require multivalent interactions

Another assumption of biomolecular condensates is that they arise from multivalent protein interactions creating a network within the dense phase^17^. However, in our interpretation of the literature, it has never been established that multivalency is required. Intriguingly, several recent findings show that phase separation can arise from very simple and short peptides^18,19^. To determine if multivalency is indeed needed for condensate formation we measured the ability of phenylalanine derivatives to form liquid-like phase separations *in vitro*. Phenylalanine with a boc protected amine (BocF) was soluble and diffuse at 75 mM in high pH where the carboxylic acid will be negatively charged (**Fig. 3A**). However, upon addition of acid to lower the pH, where the molecule will be charge neutral, BocF formed liquid-like phase separation that showed classic condensate behavior such as merging (**Fig. S3A**). Interestingly, BocF dissolved directly in low pH was insoluble above 10 mM in our hands.

**Figure 3:**
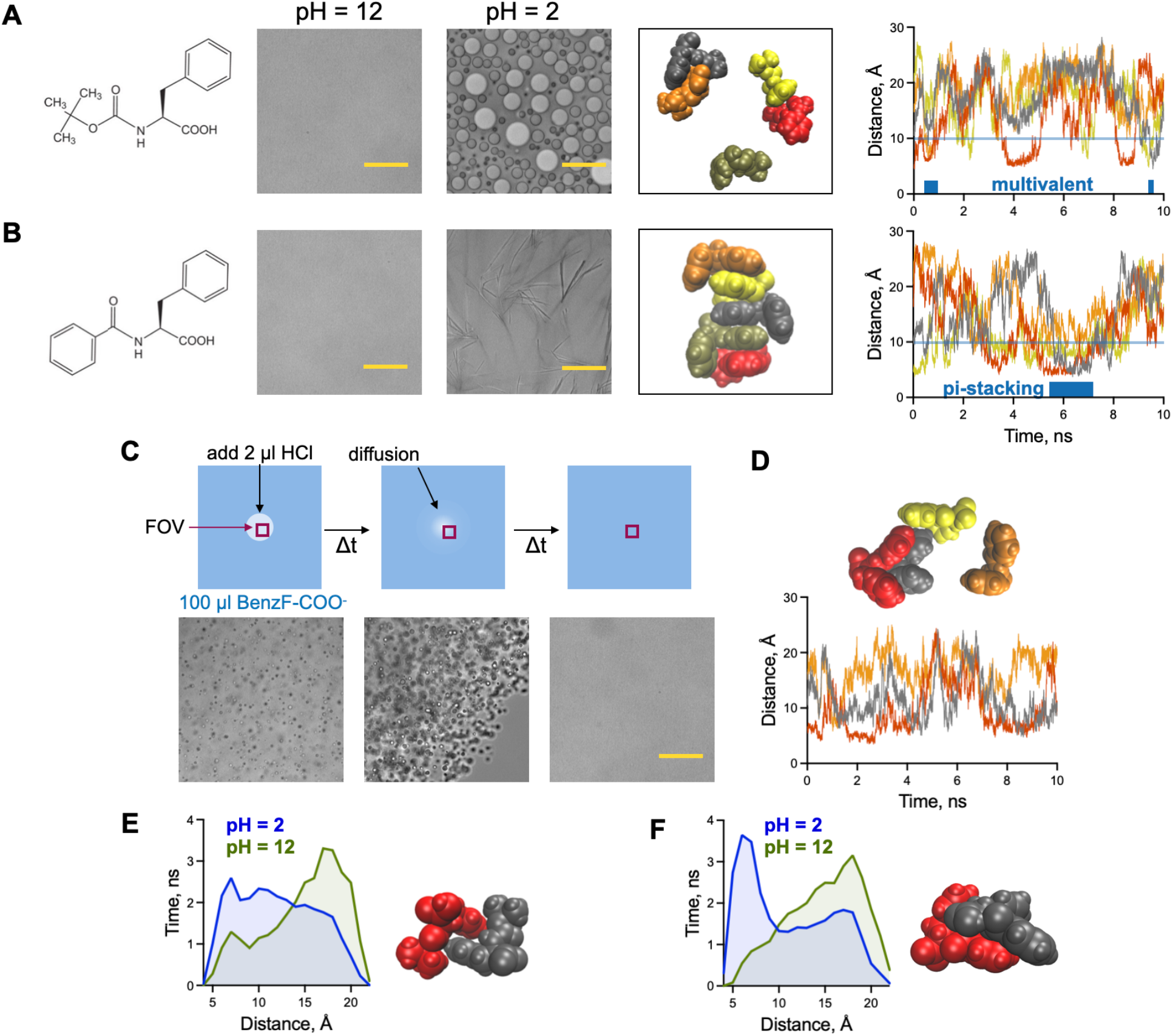
Phase separation of single phenylalanine derivatives. **A)** Boc phenylalanine is diffuse at 75 mM in pH 12 solution but forms liquid-like phase separation when the pH is lowered to 2 with the total concentration at 75 mM, shown are representative DIC images. Molecular dynamics simulations of five BocF molecules reveal the distribution of the molecules over 10 nanoseconds. The green molecule undergoes multivalent interactions with the other molecules for short durations, spending the majority of the simulation interaction with one or no other molecules. Shown is the center of mass distance between the green molecule and each of the other molecules. **B)** Benzoyl phenylalanine is also diffuse at 75 mM and ph 12, but forms solid, fibril-like structures at 75 mM when the pH is lowered to 2. Molecular simulations of BenzF molecules demonstrate that a pi-stacked, multivalent orientation can be formed, but is transient. **C)** Addition of acid to 75 mM BenzF at pH 12 generates an opaque spot that is comprised of liquid-like phase separation. The localized region of phase separation quickly disappears as the acid diffuses and is buffered return the field of view (FOV) to diffuse distribution at pH 12. **D)** Molecular dynamic simulation of two neutral BenzF and two deprotonated BenzF suggests that pi-stacking does not occur and the molecules undergo no, dimer and multivalent interactions. **E)** Molecular simulation results of two BocF molecules at pH 2 (charge neutral) and pH 12. Shown are the duration of the center of mass distance in 1 angstrom bins integrated over three separate simulations and an example structure of the two neutral molecules interacting. **F)** Molecular simulation results of two BenzF molecules at pH 2 (charge neutral) and pH 12. Shown are the duration of the center of mass distance in 1 angstrom bins integrated over three separate simulations and an example structure of the two neutral molecules interacting. All scale bars = 30 μm.

The small size of BocF suggests it cannot undergo multivalent interactions in a similar manner as long proteins or peptides. To explore if BocF forms multivalent interactions we performed molecular dynamic simulations of multiple neutral BocF molecules in Amber. A ten nanosecond simulation showed that, although multiple BocF molecules could interact, the molecules did not form a multivalent network and BocF exists as a dimer or monomer for the majority of the simulation (**Fig. 3A**). Here, measured distance between a single molecule and each of the other molecules revealed the low frequency at which more than two molecules are within 1 nm of each other. Another derivative, benzoyl phenylalanine (BenzF), where the primary amine is protected by a benzoyl group introducing a second phenyl ring, was also diffuse at high pH but when the pH was lowered formed fibril like structures (**Fig. 3B**). Molecular simulations of multiple neutral BenzF molecules found that BenzF is capable of undergoing pi-stacking, suggesting that fibril formation creates geometry where each molecule is interacting with two other molecules, generating dense interactions that are expected in solid phase separation. However, the pi-stacked formation in our simple simulation was transient, and molecules failed to form a multivalent network.

The transient nature of the BenzF simulation suggested that BenzF could form condensates under the right conditions. BenzF in stable pH at multiple pH values failed to generate condensates. However, when we added a small drop of acid to locally decrease the pH of 75 mM BenzF in pH 12, transient condensates were generated (**Fig. 3C**). The condensates rapidly dissolved and the pH returned to 12 as the acid diffused and was buffered. We considered that the dynamics of protonation and deprotonation of the carboxylic acid are more rapid in unstable acidic conditions than diffusion of BenzF molecules and ran a simulation with a mixture of charged and neutral BenzF (**Fig. 3D**). No pi-stacking was observed, suggesting that transient charge prevents more ordered structures. Furthermore, although interactions with more the one molecule occur, formation of a network of interactions was absent, suggesting that separation occurs through simple interactions. We observed a similar response of BocF to local addition of acid in a high pH solution (**Fig. S3B**).

The size of BocF and BenzF and the molecular simulation results suggest that molecules are not undergoing multivalent interactions. Furthermore, simulation results show that neutral molecules do not form hydrophobic separations. BocF was soluble up to 200 mM in pH 12 but formed solid phase separation upon lowering the pH to 2 when the resulting concentration was above 100 mM, further demonstrating that liquid-like phase separation is not purely hydrophobic in nature, and thus unlike oil in water. To determine if non-multivalent (dimer) interactions were elevated when liquid-like phase separation occurred we measured the distance between two BocF molecules over a simulation. Here, canonical measurements of binding energy and affinity are not possible as the bound structure is unknown and a BocF dimer likely has multiple lower energy interactions that could represent the bound state. Therefore, we calculated the distance between two molecules and integrated the total time spent at each distance, in 1 angstrom bins, over three separate ten nanosecond simulations. The simulation results show that two charge neutral BocF molecules spend a longer duration closely interacting with each other than charged BocF molecules, which are present at high pH (**Fig. 3E**). Simulations with two BenzF molecules found similar results with more time spent in close proximity when the molecules are in a charge neutral state (**Fig. 3F**). The shape of the neutral BenzF distance versus time histogram suggests molecules were interacting when the center of mass distance between them was below 1 nm. Therefore, we interpret a longer duration in close proximity as representing a stronger interaction affinity. Both neutral BenzF and neutral BocF have a higher dimer affinity than their charged counterparts, as would be expected. And, neutral BenzF dimer interactions had a higher affinity than neutral BocF dimer interactions (**Fig. S3C**).

Lastly, BocF with a fluorine atom on the phenyl ring (fluorBocF) also generated condensates at low pH (**Fig. S3D**). Both BocF and fluorBocF showed lower condensate abundance at decreasing concentrations. Yet, fluorBocF formed condensates at 10 mM while BocF did not, demonstrating the fluorBocF phase separation has a lower C_sat_.

Molecular simulations of fluorBocF dimer interactions found that the fluorinated form had a slightly higher dimer affinity than BocF (**Fig. S3E**), revealing that dimer binding affinity correlates with C_sat_.

To determine if monovalent interactions could drive condensation of peptides we designed 10mer peptides as polylysine and polyglycine block copolymers with different polylysine lengths. We had previously found that polylysine mixed with RNA forms aggregates^20^. However, here we reduced the pH of the solution and found that 10mer polylysine mixed with RNA forms condensates at pH 3 (**Fig. S4A**). We then determined the ability of the copolymers to form condensates with RNA. We found that condensates formed with peptides having polylysine lengths down to 4 amino acids of the 10mer at pH 3, but no phase separation was observed with only 2 lysines in the 10mer. We confirmed that RNA is necessary to form condensates by testing the 10mer amino acid with 6 lysines in the absence of RNA and observed no phase separation (**Fig. S4B**). These results demonstrate that positively charged lysine peptides can form condensates with RNA and only four lysines are necessary in our system. The limited length of 4 lysines suggests that multivalency is not possible in these phase separations and supports the thesis that multivalency in not necessary for condensate formation.

## Discussion

Liquid-liquid phase separation describes the thermodynamics of phase separation through a phase diagram that spans a dilute system, phase separation range and a dense system (**Fig. 1A**). LLPS defines the partition coefficient as independent of total concentration, but here we demonstrated that phase separation in cells generated a partition coefficient that was dependent on the total concentration. Previous analysis of this phenomenon had assigned this concentration dependency to the presence of other species within a condensate leading to heterotypic interactions. However, when we analyzed condensation of FUS-GFP *in vitro*, a pure homotypic condensing protein, we found that the partition coefficient is concentration dependent, indicating that condensates are not LLPS. The lack of impact from heterotypic interactions is further supported by our observation that measurements from four cell lines fell along a single curve, suggesting that differences between these cell lines do not impact the partition coefficient dependency on concentration. Overall, four different proteins, from measurements *in vitro* and in cells, display a similar concentration dependent partition coefficient.

The long, disordered length of many condensing proteins suggests that multivalent interactions are abundant and therefore must be involved in condensate formation. However, this has not been explicitly shown. Our findings that phenylalanine derivatives form liquid-like phase separation demonstrates that multivalency is not required for phase separation. Their small size dictates that to form a multivalent network these molecules would have to exist at a very high concentration in the dense phase, greater than 1 M. However, BocF at a pH where it would have no charge was not soluble above 100 mM, therefore the dense phase simply cannot reach a concentration where multivalency exists. Indeed, BenzF simulation results can form multivalent interactions, but only at very high concentrations that generate solid, not liquid, phase separation. Furthermore, BocF in a liquid like phase separation is not undergoing immiscible separations as the dense phase is liquid-like and BocF is a solid at concentrations where it is not soluble. Although phenylalanine derivatives are not as long as phase separating peptides and proteins, there is no reason why they would not be governed by the same thermodynamic principles.

Accepting that biological phase separation is not driven by traditional LLPS physics or multivalent networks, a new model for how condensates arise is needed. We recently introduced the dimer model of phase separation^20^ that proposes condensates arise through rapid dimer binding interactions (**Fig. 4**). The dimer model predicts that biomolecular condensation creates an energetic minimum through increased dimer binding interactions at higher condensate concentrations. Here, at certain concentrations and binding affinities increased dimerization at higher dense phase concentration overcomes the entropic cost of phase separation to establish condensates as the minimum energetic state. The dimer model predicts that only monovalent binding interactions are necessary to form condensates, congruent with our molecular simulations of phenylalanine derivatives. Furthermore, the model predicts a total concentration dependent partition coefficient where the partition coefficient decreases with increasing concentration, a prediction that mirrors the experimental results found here.

**Figure 4.**
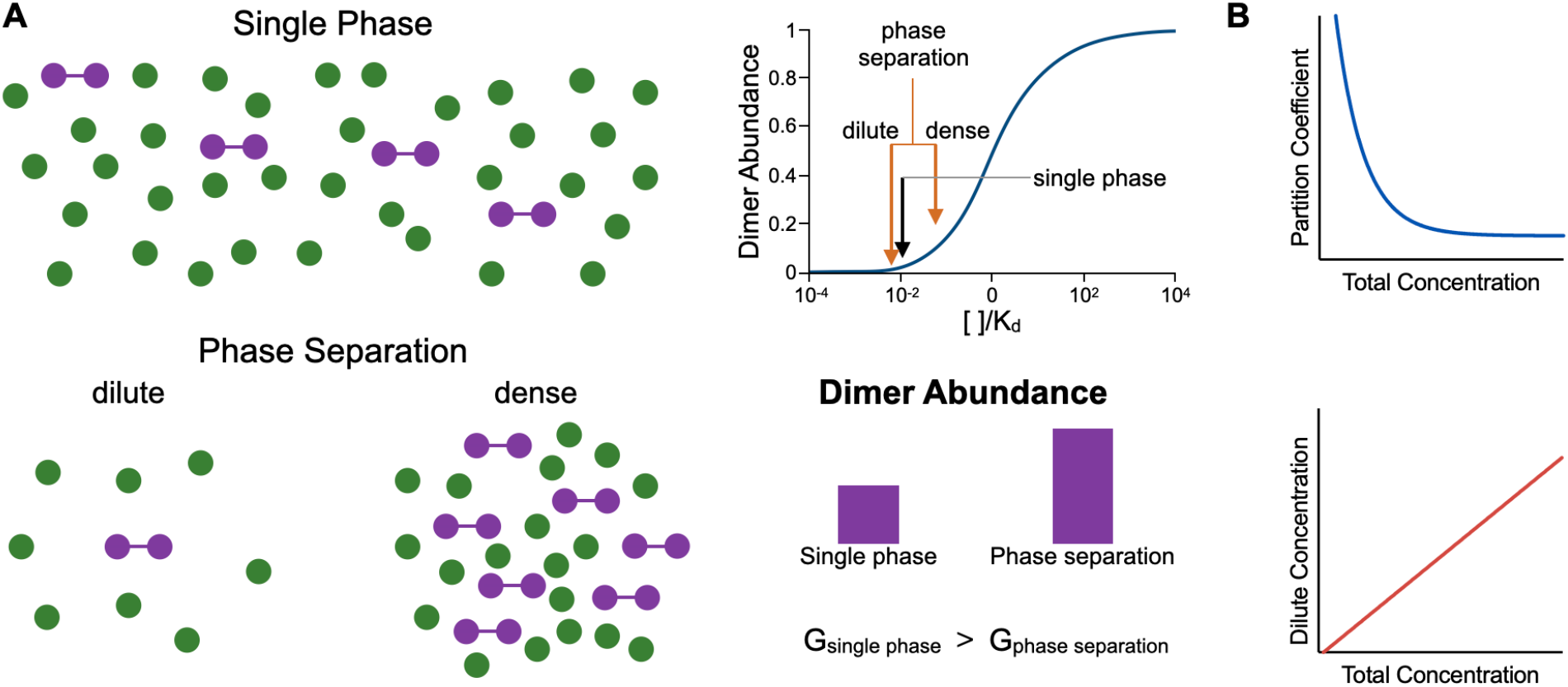
Dimer interactions in phase separation. **A)** In a single phase the dimer abundance is dictated by the concentration and affinity. Under phase separation where a dense phase is created the dimer abundance is a sum of both the dimers in the dilute phase and in the dense phase. The total abundance of dimers can be higher under phase separation, which creates a lower Gibbs free energy of binding of the system. **B)** The dimer model of phase separation predicts a partition coefficient and dilute phase concentration dependence on the total concentration that is very similar to experimental results.

Overall, results here demonstrate that liquid-like phase separation is not classic liquid-liquid phase separation, an important distinction when considering the physics driving phase separation, and that multivalent networking interactions are not required. We expect that clarifying these misconceptions of biomolecular condensate formation will have tremendous impact in understanding the thermodynamics of phase separation in biology. The demonstration that condensates can arise from monovalent interactions and have concentration dependent partition coefficients suggests that models such as the dimer model may have merit.

## Methods

### Cell Culture

The human lung cancer line A549, the human breast cancer cell line MCF7, the human ovarian cancer cell line OV90, and the triple negative breast cancer cell line HCC1937, and were cultured with Roswell Park Memorial Institute (RPMI) media supplemented with fetal bovine serum (FBS) and penicillin/streptomycin. Cells were cultured at 37 °C in a 5% CO_2_ atmosphere. A549, MCF7, OV90, and HCC1937 cells were seeded at a density of 3000 cells per well in a 384 well plate and incubated overnight to allow cells to adhere. To generate G3BP1 intracellular condensate formation cells were treated with 500 μM or 1 mM sodium arsenite. Cells were treated for one hour before fixation.

### Immunofluorescence

Cells were fixed with 4% PFA in PBS for fifteen minutes at room temperature. Cells were then washed with 1X PBS twice. 25 μL of Blocking Buffer comprised of 5% normal goat serum (CST #5425) and 0.3% Triton X-100 in 1X PBS was added to the plate and rocked for one hour at room temperature. Primary antibody was diluted 1:500 in antibody dilution buffer comprised of 1% BSA and 0.3% Triton X-100 in 1X PBS. Cells were incubated on a rocker overnight at 4 °C. Cells were then washed three times with 1X PBS and incubated with secondary antibody, anti-mouse IgG Fab2 Alexa Flour (R) 555 (1:1000 in antibody buffer) for one hour on rocker at room temperature and protected from light. The cells were then washed three times with 1X PBS and stained with DAPI (1 μg/ml in 1X PBS) for five minutes. Finally, the plate was washed twice more with 1X PBS. Imaging was performed on a Nikon Ti-2 fluorescent microscope with a 20x air objective.

### Cellular Analysis

All analysis was performed in Python. For each immunofluorescence image, background noise, determined as the fifth percentile of total image intensity, was subtracted from the fluorescence channel. For PML and NPM1, which produce nuclear phase separation, single cell nuclei were segmented using the Stardist cellular segmentation package on the DAPI fluorescence image. Nuclei were filtered by size to remove any poor segmentations. Sub-nuclear segmentation of dense and dilute phases was done via multi-Otsu thresholding of intensity values with four regions. The highest threshold values represent the condensates. The second most intense threshold region was omitted to avoid any unclear regions and areas were scattered light from the condensates increases intensity in the image. The third threshold identified the dilute phase or non-condensate region, and the fourth Otsu threshold region was the near-zero background as well as nucleoli in the case of PML (**Fig. S1A**).

For cytoplasmic condensate images, cells were segmented with the Cellpose segmentation package to generate masks for each image. Data extraction was done using the regionprops table function from the sklearn package. To isolate cytoplasmic regions, we binarized and inverted the nuclear masks and multiplied this array by the cytoplasm masks, effectively removing each cell nucleus. To remove poorly segmented cells, we used the cytoplasm masks to identify corresponding nuclear masks and removed all cells lacking exactly one nucleus. Similar to the approach used for nuclear condensate images, the segmentation of condensates and the dilute phase was done via multi-Otsu thresholding intensity values with four regions. The values in the highest threshold delineated condensates, followed by an excluded region surrounding condensates, accounting for light diffraction effects and unclear regions. The third threshold was the dilute phase/non-condensate region, and the fourth was the near-zero background as well as the removed nuclear region (**Fig. S1A**).

Following the segmentation of all cells into distinct regions, the partition coefficient for each cell was calculated as the ratio of the average intensity of the dense phase region to that of the dilute phase region. To ensure the inclusion of only cells containing condensates in our dataset, we filtered out cells based on the average intensity of the dense region. The filtering threshold value was determined by comparing two images: one with clearly visible condensates and one without. Once filtered to include only cells where the average intensity was above a range between 500 and 2,500 (arbitrary units), we found that all cells with condensates were retained while 94.5% of cells without condensates were removed when the threshold was set to 1,000 au (**Fig. S1B**). Thus, our data was filtered across all cell lines to include only cells where the dense phase condensate region average exceeded 1,000 au (**Fig. S1B**).

### FUS-GFP phase separation

pET MBP GFP LIC cloning vector was a gift from Scott Gradia (Addgene #29750), MBP-FUS_FL_WT was a gift from Nicolas Fawzi (Addgene #98651). FUS was cloned using the plasmids (forward TACTTCCAATCCAATGCAATGGCCTCAAACGATTATACCC, and reverse CTCCCACTACCAATGCCATACGGCCTCTCCCTG) by PCR and gel purified. The LIC cloning vector was digested by SspI (NEB #R3132) and gel purified. The DNA was then digested with LIC qualified T4 polymerase (Sigma #70099), annealed following the LIC cloning protocol and transformed into BL21 (NEB #C2527) cells. This ligation created a plasmid that expresses a single fused protein containing: his-tag, maltose binding protein, TEV cut site, FUS and GFP under a lac operator. The plasmid was confirmed by whole plasmid sequencing (MGH DNA Core).

BL21 cells were grown at 37 °C in LB broth with kanamycin until they reached an OD of 0.6. 500 μM of IPTG was then added and cells were cultured at 37 °C for 3 hours. The cells were then pelleted and resuspended in protein solution (in 1 M KCl, 1 mM DTT, 50 mM Tris,10 mM imidazole, and 5% glycerol pH 7.4. Cells were snap frozen in liquid nitrogen, thawed and disrupted by probe tip sonication on ice, 30 seconds on 30 seconds off for 10 minutes. The solution was cleared by centrifugation at 20,000g for 30 minutes. The supernatant was the incubated at 4 °C with 2 ml of washed Ni-NTA agarose beads (Qiagen) for 1 hour. Beads were washed 3 times with 25 ml of protein solution in a gravity column and the protein was eluted with protein solution containing 300 mM imidazole. Protein was dialyzed in protein solution in a 3 kDa dialyzer (Pierce) to remove excess imidazole. FUS-GFP was cleaved from MBP using TEV protease with a His tag (NEB) at 30 °C for 2 hours and collected as wash through on Ni-NTA beads. FUS-GFP concentration was determined by protein absorbance at 280 nm and verified by GFP absorbance at 504 nm. Protein was snap frozen and stored at -80 °C.

To form FUS-GFP condensates, protein was diluted in solutions to reduce the concentration of KCl and glycerol to 150 mM and 0.5% for all FUS-GFP concentrations measured. Diluted protein solution was imaged in glass bottom 96 well plates (CellVis) with a 60x NA = 1.4 oil immersion objective on a Nikon Ti-2 fluorescent microscope.

### Phenylalanine phase separation

Phenylalanine and derivates (Chem-Impex) were dissolved in solution containing 10 mM Tris and 100 mM NaCl at the desired pH. Compounds were diluted to the desired concentration and pH. Solutions were immediately transferred to a 96 well plate mounted on the microscope and imaged with DIC using a 60x NA 1.4 oil immersion objective. Transient formation of phase separation was achieved by adding 2 μl of 6 M HCl to compounds in Tris/NaCl buffer at a pH of 12.

### Peptide-RNA phase separation

Peptides (Genscript) and RNA (IDT) were dissolved in solution containing 10 mM Tris and 100 mM NaCl, with a pH of 5 or 3. Solutions were mixed to obtain the desired concentrations, immediately transferred to a 96 well plate mounted on the microscope and imaged with DIC using a 60x NA 1.4 oil immersion objective.

### Molecular Dynamic Simulations

Simulations were run in Amber using the gaff2 force field for phenylalanine derivates and the OPC force field for water. Phenylalanine derivative structures were generated in Flare using the chemical SMILES. The structures were saved in mol2 format and optimized for Amber simulations using antechamber. Molecules were solvated with explicit water and an 8.0 Å buffer distance in Leap. Energy was minimized at constant pressure and the molecules were heated in Sander. Simulations were carried out with pmemd at constant pressure. Data were acquired every 2 fs. Results were visualized and rendered in VMD and distances were measured using cpptraj.

