## Supplementary Figures for "Clarifying misconceptions of biomolecular condensate formation"

**Affiliations:**
Institute for Innovation in Imaging, Department of Radiology, Massachusetts General Hospital, Harvard Medical School, Boston, MA

**Correspondence:**


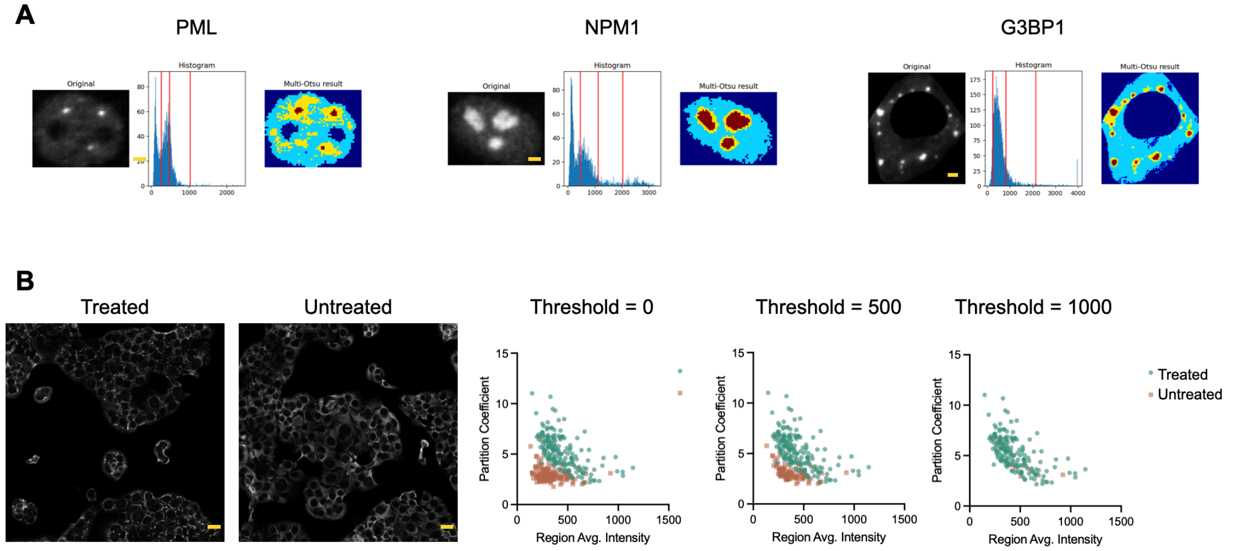


**Figure S1: Method to determine intracellular partition coefficient. A)** Example images of a single A549 nuclei or cell labeled for PML, NPM1 or G3BP1. Shown are the corresponding multi-Otsu thresholding histogram and graphical depiction of the classes assigned by three thresholds. Shown are pixels from each intensity class ranging from highest intensity (red) to lowest (dark blue). Here the lowest intensity class lacks fluorescent signal. The higher middle class (yellow) is ignored because these pixels are most difficult to classify and the partition coefficient is calculated from the average intensity of red pixels divided by the average intensity of light blue pixels. Scale bars are 10 µm for PML and NPM1 and 10 µm for G3BP1. **B)** Representative images of A549 cells with G3BP1 immunofluorescence that were untreated or treated with arsenite. Untreated cells show few condensate foci. Plotted are the calculated single cell partition coefficients as a function of total region intensity for the two images (threshold = 0). A highest intensity average value threshold of 500 au selects for cells that have G3BP1 condensates and removes some of the untreated cells from the measurements but does not impact the treated cells. Increasing the threshold to 1,000 au nearly removes all untreated cells without removing any treated cells. An intensity threshold of 1,000 was applied to all images to remove erroneous partition coefficient calculations from cells that do not have phase separation.  Scale bar = 10 µm.

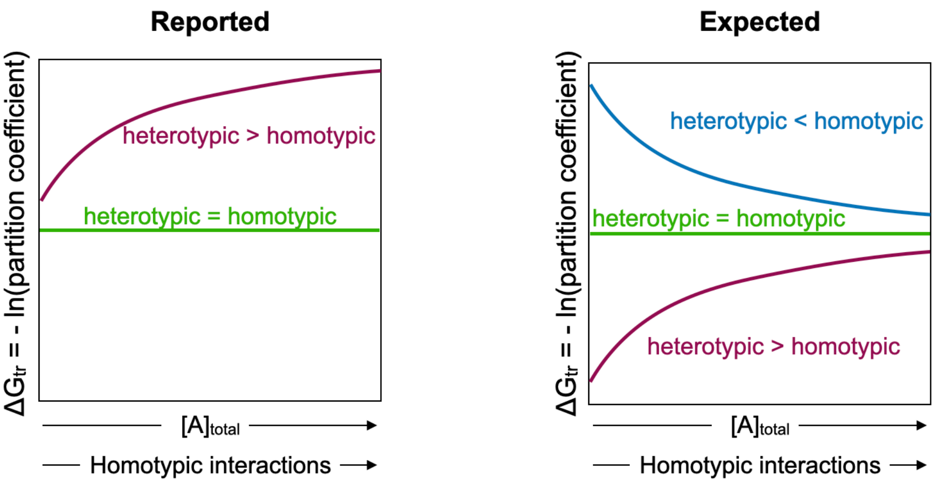


**Figure S2: Impact of heterotypic interactions on the free energy of transfer. Left**, reported (Riback, *et al*)[14] energy for a system with two components as a function of increasing the concentration of one component, A. When the interaction energy is the same, the energy is independent of the relative abundance of each component (green). The equal interaction energy curve is the same as expected for single component systems. When heterotypic interactions generate a lower energy the total energy increases with increasing homotypic interactions, or increasing concentration of a single component (purple). **Right**, the expected energy dependence on homotypic interactions. When heterotypic interactions are more energetically favorable the system energy is lower than a single component system. Also, as homotypic interactions increase for both heterotypic > homotypic and heterotypic < homotypic the total energy converges with a pure, single component system as homotypic interactions increase.

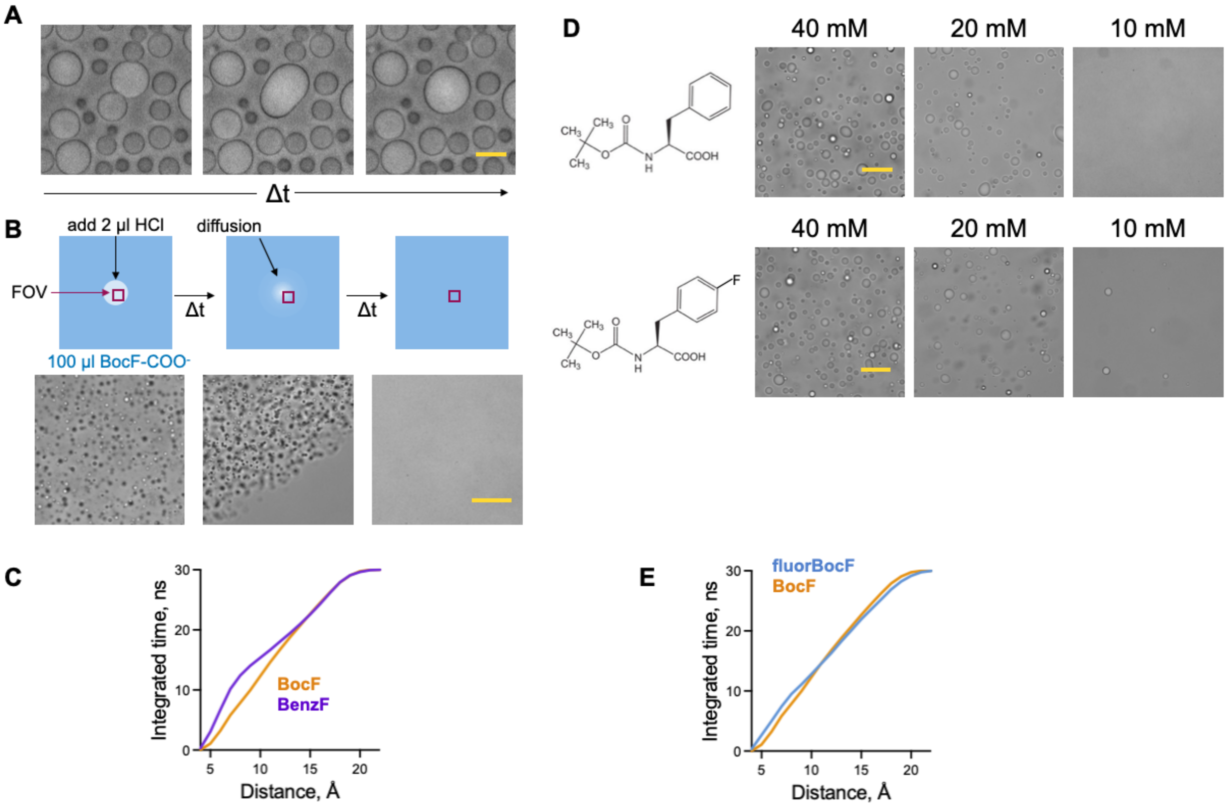


**Figure S3: Phase separation of phenylalanine derivatives. A)** Representative images of BocF condensates merging over 2 seconds, each frame is one second time difference, scale bar = 10 µm. **B)** BocF transient condensate formation following local addition of acid, scale bar = 30 µm. **C)** Molecular simulation results for two molecules of BocF (orange) or BenzF (purple) at pH 2. Plotted are running sums of the duration of the distance between the two molecules, binned at 1 angstrom. Here, BenzF spends are larger amount of time at shorter distances than BocF. **D)** Phase separation of BocF and fluorBocF show different concentrations at which condensates arise. Shown are representative images, scale bar = 30 µm. **E)** Running integration of the duration of the distance between two molecules of BocF (orange) or fluorBocF (blue) at pH 2 from simulation results. FluorBocF has a slightly longer duration of close range interactions.

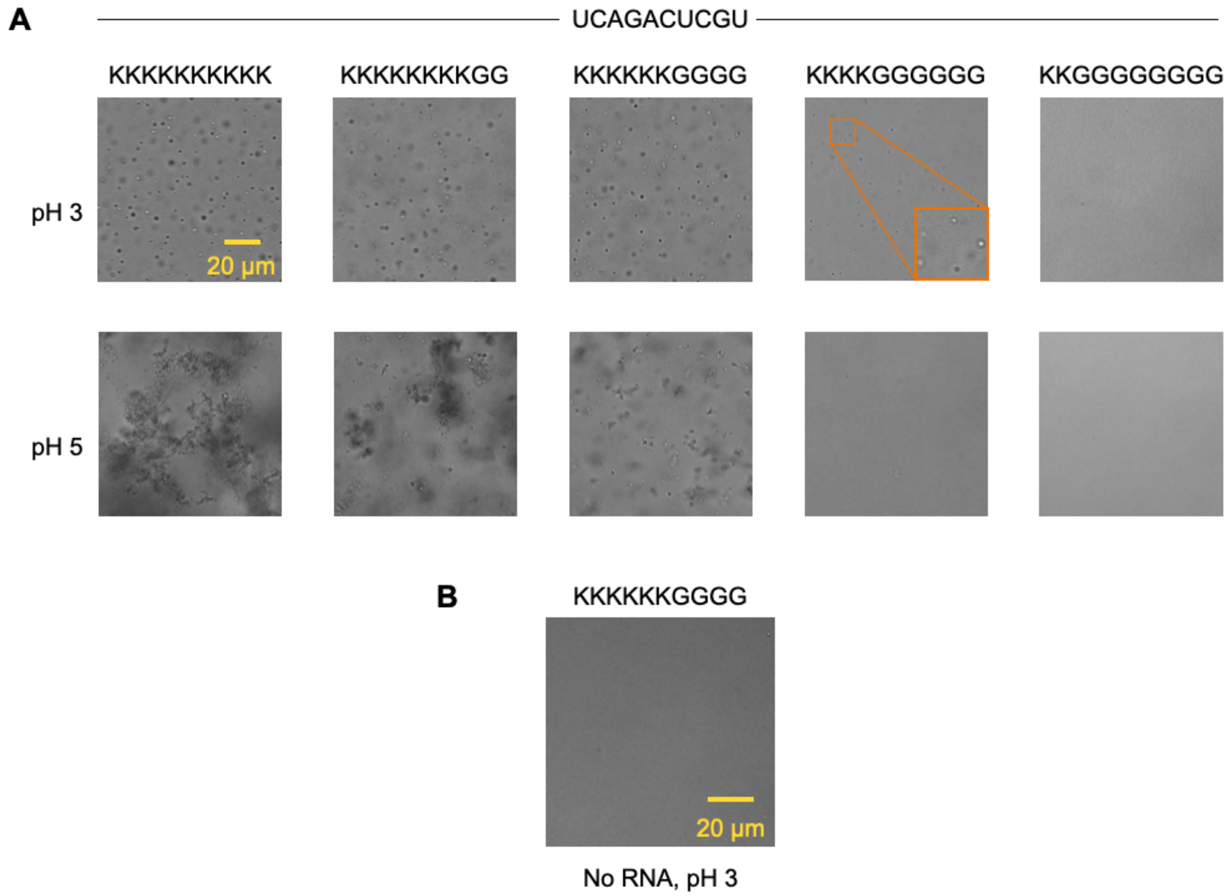


**Figure S4: RNA-peptide phase separation. A)** Representative images of a 10mer RNA mixed with a 10mer peptide at pH 3 or pH 5. Liquid like phase separation occurs at pH 3 with only 4 lysines in the peptide. **B)** Representative image of a peptide in the absence of RNA. All concentrations 10 mM.
